# Slow gamma oscillations in the mouse olfactory bulb are correlated with sniffing in the dark period

**DOI:** 10.1101/2023.04.25.538246

**Authors:** Ryosuke Mochizuki-Koike, Mami Okada, Yuji Ikegaya, Nobuyoshi Matsumoto

## Abstract

Neural activity in the olfactory bulb is reflected in local field potentials (LFPs). Functionally, LFPs in the olfactory bulb are categorized into different frequency bands: 1-4 Hz, 6-12 Hz, 25-50 Hz, and 65-130 Hz, which respectively correspond to respiration, sniffing, slow gamma, and fast gamma oscillations. While gamma oscillations in the olfactory bulb are modulated by respiration and sniffing, it remains unknown how and whether the modulation of LFP oscillations is affected by the time of day. To address this question, we recorded LFPs in the olfactory bulb, hippocampus, and neocortex of unrestrained mice for up to 3 d. For each recording site, we calculated the correlation coefficients of normalized LFP powers between pairs of frequency bands in the three regions during the dark and light periods. We then compared these correlations with those generated by surrogate data to investigate whether the correlation was statistically significant. We found that the correlation between sniffing and slow gamma oscillations was higher in the dark period than in the light period. Our finding has the potential to shed light on the coding scheme of olfactory information that is dependent on the light/dark cycle.

## 1. Introduction

The olfactory system plays a crucial role in detecting and perceiving odors for food selection, predator avoidance, and social communication [1–3]. The odor information processing is supported by neural activity in the olfactory bulb, the first relay station in the olfactory system [4]. This olfactory neural activity is electrophysiologically reflected by extracellular oscillations called local field potentials (LFPs) [5].

LFPs are often bandpass-filtered at a certain frequency range to investigate frequency band-specific functions relevant to behavior. For example, the relatively low-frequency component (*e*.*g*., 1-12 Hz) of LFPs in the olfactory bulb represents inspiratory-expiratory cycles in association with nasal airflow [4–10]. This nasal breathing pattern is further classified into slower (<4 Hz) and faster (>6 Hz) components that usually indicate respiration and (fast) sniffing, respectively. In particular, (fast) sniffing is involved in active perception that contributes to motivation for exploration and social interactions [11,12]. In contrast, the high-frequency component (25-100 Hz) of LFPs is called gamma oscillations produced in various regions, including the olfactory bulb, hippocampus, and neocortex. While gamma oscillations in the hippocampus and neocortex are dependent on neural network activity created by the interaction of inhibitory interneurons alone and/or the interaction between excitatory pyramidal cells and interneurons (*e*.*g*., fast-spiking parvalbumin-positive basket cells) [13,14], the interaction of excitatory mitral cells and inhibitory granule cells at the dendrodendritic reciprocal synapses is believed to contribute to the generation of gamma oscillations in the olfactory bulb [14,15]. Since the neural activity of mitral cells could be modulated by peripheral input (*i*.*e*., odorants and mechanical stimuli caused by nasal airflow) from olfactory sensory neurons in the nasal epithelium [16,17], the gamma oscillations are indeed affected by respiration [15]. Consistent with the previous study [15], when gamma oscillations are classified into two types of oscillations with different frequency subbands (*i*.*e*., slow and fast), the slow and fast gamma oscillations in the olfactory bulb could be contributed by the activity of mitral and granule cells, respectively, and paced by sniffing rhythm [18]. These studies suggest that the respiratory/sniffing components of the olfactory bulb LFPs are tightly coupled with the gamma oscillations; however, no studies have scrutinized how the correlation between the frequency band-specific (*i*.*e*., respiration, sniffing, slow gamma, and fast gamma) components of the olfactory bulb LFPs dynamically changes throughout the day.

To address this question, we simultaneously recorded LFPs in various sites of the olfactory bulb as well as the hippocampus and neocortex of freely behaving mice for up to 3 d and bandpassed the long-term LFPs at the specific frequency range indicative of respiration, sniffing, slow gamma, and fast gamma for the olfactory bulb, and delta, theta, slow gamma, and fast gamma for the hippocampus and neocortex. We then divided the bandpassed LFPs into the dark and light periods and calculated correlation of the bandpassed LFPs between all pairs of frequency bands for all recording sites.

## 2. Materials and methods

### 2.1 Ethical approval

Animal experiments were performed with the approval of the Animal Experiment Ethics Committee at The University of Tokyo (approval number: P29-14) and according to the University of Tokyo guidelines for the care and use of laboratory animals. These experimental protocols were carried out in accordance with the Fundamental Guidelines for the Proper Conduct of Animal Experiments and Related Activities in Academic Research Institutions (Ministry of Education, Culture, Sports, Science and Technology, Notice No. 71 of 2006), the Standards for Breeding and Housing of and Pain Alleviation for Experimental Animals (Ministry of the Environment, Notice No. 88 of 2006) and the Guidelines on the Method of Animal Disposal (Prime Minister’s Office, Notice No. 40 of 1995). All efforts were made to minimize animal suffering.

### 2.2 Animals

A total of five male 9-to 12-week-old C57BL/6J mice (Japan SLC, Shizuoka, Japan) were housed under conditions of controlled temperature and humidity (22 ± 1 °C, 55 ± 5%) and maintained on a 12:12-h light/dark cycle (lights on from 7:00 to 19:00) with *ad libitum* access to food and water.

### 2.3 Preparation

A recording interface assembly was prepared as previously described [19,20]. In short, multiple tetrodes, each of which was constructed by bundling together four 17 μm polyimide-coated platinum–iridium alloy (90/10%) wires (California Fine Wire, USA), were prepared. The depth of the tetrodes from the brain surface was independently adjustable. The electrode assembly was composed of an electrical interface board (EIB) (EIB-36-PTB, Neuralynx, USA) and custom-made bodies by three-dimensional (3-D) printers (Formlabs Form 2, Formlabs, USA). The EIB had a sequence of metal holes for connections with the alloy wires. A given individual hole was conductively connected with one end of the insulated wire using attachment pins, whereas the opposite end was the tip of tetrodes. The platinum coating was applied to the tip of each tetrode so that the impedances of the tips would range from 150 to 300 kΩ measured with an impedance tester (nanoZ, White Matter, USA).

### 2.4 Surgery

For the mice surgery, general anesthesia was induced and maintained with 3% and 1.0-1.5% isoflurane gas, with careful inspection of the animal’s condition during the whole surgical procedure. The veterinary ointment was applied to the mouse’s eyes to prevent drying. The skin was sterilized with povidone-iodine and 70% ethanol whenever an incision was made.

After anesthesia, a mouse was mounted onto a stereotaxic apparatus (SR-6M-HT, Narishige, Japan). The scalp was incised with a surgical knife. A circular craniotomy with a diameter of approximately 0.9 mm was performed using a high-speed dental drill (SD-102, Narishige). Two stainless-steel screws (0.8 mm in diameter, 3 mm in length) were implanted into the bone above the cerebellum as ground and reference electrodes as previously described [19,21]. An epidural screw electrode was stereotaxically implanted into the olfactory bulb to record LFPs, whereas tetrodes were implanted into the neocortex and hippocampus. The exposed brain surface around the tetrodes was covered using a silicone elastomer (Kwik-Sil, World Precision Instruments, USA). The exposed skull was then covered using a dental resin cement (Super-Bond C&B Kit, Sun Medical, Japan). The tetrode assembly was secured on the skull using an acrylic resin cement (Re-fine Bright, Yamahachi Dental, Japan). A wire electrode (AS633, Cooner Wire, USA) was implanted into the trapezius to record electromyograms. The screw and wire electrodes were connected with the EIB using conductive wires and covered using acrylic resin cement. The electrode assembly was protected by a custom-made hood.

While our experimental protocols mandate the humane killing of animals if they exhibit any signs of pain, prominent lethargy, or discomfort, such symptoms were not observed in any of the five mice used in this study.

### 2.5 *In vivo* electrophysiology

Following surgery, each mouse was allowed to fully recover from anesthesia and was housed individually in transparent plexiglass cages with free access to water and food. The tetrodes were lowered by a step of approximately 250 μm to reach 1,000 μm under the brain surface. Then, the tetrodes were carefully lowered by approximately 31.25-62.5 μm until hippocampal ripples with large amplitudes were observed.

Neural signals and electromyograms were recorded in the home cage (with free access to water and food) for up to 3 d using a data acquisition system (CerePlex Direct, Blackrock, USA). The electrophysiological signals were digitized at 2 kHz and lowpassed at 500 Hz.

### 2.6 Histology

After the recordings, the mice were anesthetized with an overdose of urethane. Electric currents (25 μA) were applied to each tetrode to make a scar. The mice were transcardially perfused with 4% paraformaldehyde (PFA) in 0.01 M phosphate-buffered saline (pH 7.4), followed by decapitation. The brains were preserved in a cold place overnight. They were further soaked overnight in 4% PFA and overnight in 20% sucrose (in 4% PFA) twice for post-fixation and cryoprotection. The brains were frozen using dry ice to preserve at - 80°C and sectioned at a thickness of 40 μm using a cryostat (CM3050S, Leica Biosystems, Germany).

Serial slices were then mounted on glass slides and stained with cresyl violet. Cresyl violet staining was performed based on a previously described procedure [22–25]. Briefly, the slices were rinsed in water, ethanol, and xylene; counterstained with cresyl violet; and coverslipped with a mounting agent (PARAmount-D, FALMA, Japan). The positions of the electrode tips were confirmed by identifying a scar. Data were excluded from the subsequent analysis if the tip position was outside the olfactory bulb, hippocampus, or neocortex. Cresyl violet-stained images were acquired using a microscope (BZ-X710, Keyence, Japan).

### 2.7 Data analysis

All data analyses were performed using custom-made MATLAB 2022b (MathWorks, USA) routines. The null hypothesis was statistically rejected when *P* < 0.05, unless otherwise specified. When multiple comparisons were required, we corrected the original significance level (*i*.*e*., 0.05) with Bonferroni correction and compared the original *P* values with the adjusted significance level.

When apparent electrical noise (caused by the physical impact of mice hitting their heads on the walls, glooming, rearing, and chewing) dominated the signals during certain periods, these periods were excluded from the subsequent analysis using the MATLAB ‘filloutliers’ function; an outlier at a time point *t* was defined as a value (at *t*) larger than 30 times of the median absolute deviation of the values between *t*-15 and *t*+15 (*i*.*e*., ± 7.5 ms from the given time point). After removing noise contamination, all LFP signals were transformed into a frequency domain using the fast Fourier transform to yield power spectra of the neural signals.

The entire measurement period was divided into two halves: the light period (7:00 a.m. - 7:00 p.m.) and dark period (7:00 p.m. - 7:00 a.m.). LFP signals in the olfactory bulb were bandpass-filtered at four frequency bands: 1-4 Hz (slow sniffing, or ‘respiration’), 6-12 Hz (fast sniffing, or ‘sniffing’), 25-50 Hz (slow gamma), and 65-130 Hz (fast gamma). The hippocampal LFP signals were also bandpassed at 1-4 Hz (delta), 6-12 Hz (theta), 25-50 Hz (slow gamma), and fast gamma (65-130 Hz), while the neocortical signals were similarly broken down to extract delta, slow gamma, and fast gamma frequency oscillations (Fig. 1).

**Figure 1.**
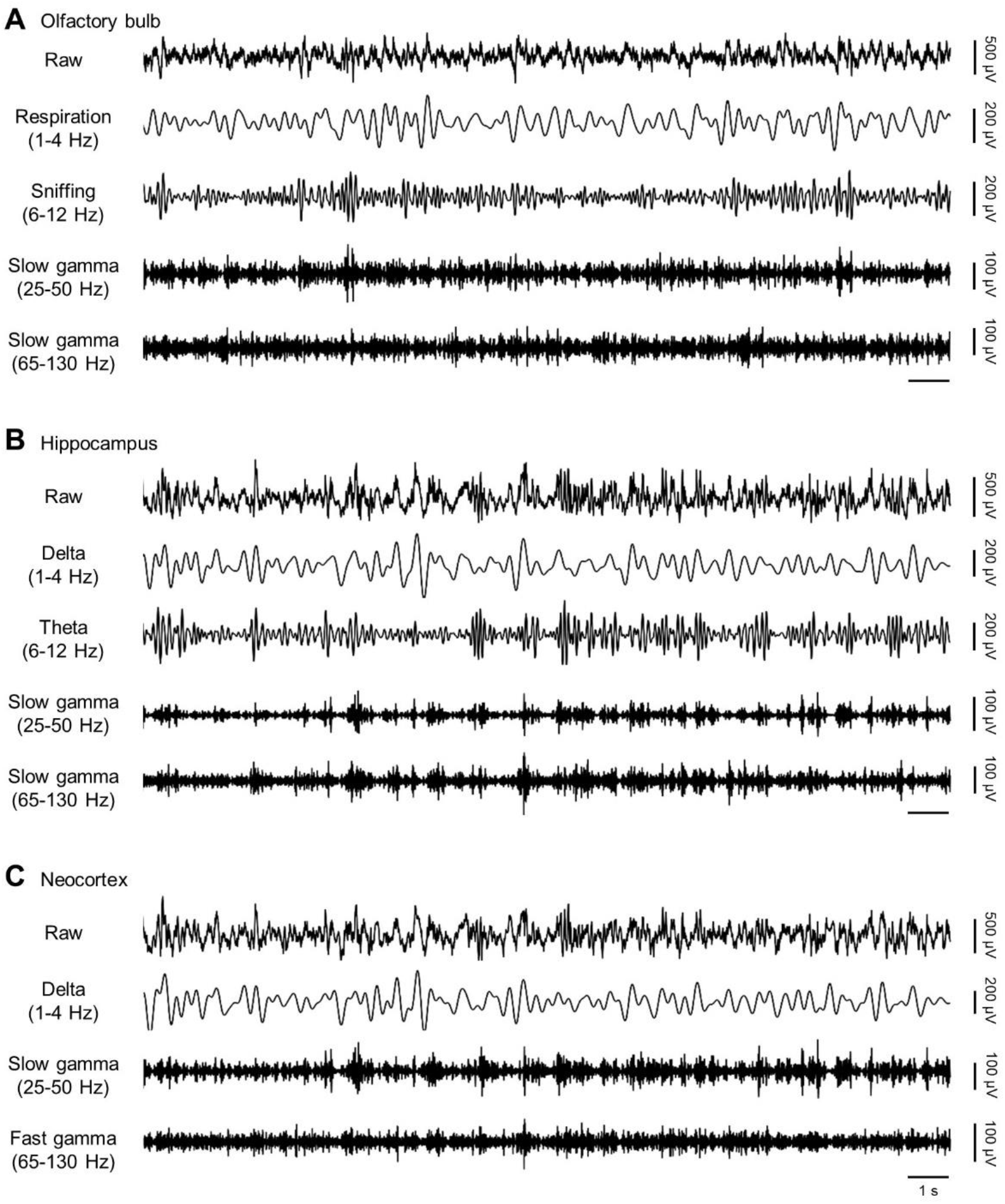
Simultaneous recordings of LFPs in the olfactory bulb, hippocampus, and neocortex. ***A***, A representative raw (*top* (*first*)) trace of LFPs in the olfactory bulb is shown. The trace is bandpassed at 1-4 Hz (*second*), 6-12 Hz (*third*), 25-50 Hz (*fourth*), and 65-130 Hz (*fifth*), each of which represents respiration, sniffing, slow gamma, and fast gamma oscillations, respectively. ***B***, A representative raw (*top* (*first*)) trace of LFPs in the hippocampus is shown. The trace is bandpassed at 1-4 Hz (*second*), 6-12 Hz (*third*), 25-50 Hz (*fourth*), and 65-130 Hz (*fifth*), each of which represents delta, theta, slow gamma, and fast gamma oscillations, respectively. ***C***, A representative raw (*top* (*first*)) trace of LFPs in the neocortex is shown. The trace is bandpassed at 1-4 Hz (*second*), 25-50 Hz (*third*), and 65-130 Hz (*fourth*), each of which represents delta, slow gamma, and fast gamma oscillations, respectively.

To calculate the correlation coefficients of the LFP power between frequency bands, the raw signals were split by 10 s. Based on the fast Fourier transform, each 10-s segment was transformed into a function of frequency to estimate the power spectrum density and calculate the power of certain frequency bands (see above) by the area under the curve. The power of certain frequency bands was then normalized into a Z-score. Correlation coefficients of the Z-scored power between all possible pairs of the frequency bands (*i*.*e*., 6 for the olfactory bulb and hippocampus, and 3 for the neocortex) were calculated. To create 10,000 surrogate correlation coefficients for each pair of the frequency bands, the original 24-h-long Z-scored power was randomly shuffled by 1 h using the Mersenne Twister generator, a pseudorandom number generator, so that a total of 24! surrogates could be theoretically generated. For each pair of frequency bands (*e*.*g*., delta *vs*. theta), a *P* value was calculated as the ratio of the number of the surrogate pairs whose correlation coefficients were higher than those of the original pair to 10,001 (*i*.*e*., surrogate data test) [26], followed by Bonferroni correction [27], which enabled us to find a ‘significantly correlated site’ for the frequency band pair. This calculation of *P* values was repeated for all cases. The number of significantly correlated sites was compared between the dark and light periods using Yates’ chi-square test.

## 3. Results

We conducted simultaneous LFP recordings in the olfactory bulb, hippocampus, and neocortex of freely behaving mice for a continuous period of 24 h. We then bandpass-filtered the olfactory LFPs with the frequency ranges of respiration (1-4 Hz), (fast) sniffing (6-12 Hz), slow gamma (25-50 Hz), and fast gamma (65-130 Hz) as well as the hippocampal and neocortical LFPs with the frequency ranges of delta (1-4 Hz), theta (6-12 Hz), slow gamma (25-50 Hz), and fast gamma (65-130 Hz) (Fig. 1).

For a given animal, we normalized each power of the bandpassed LFPs in the olfactory bulb, hippocampus, and neocortex to see if the bandpassed components in each region were chronologically correlated (Fig. 2A-C). We then divided the normalized power of each frequency band into light and dark periods and calculated correlation coefficients for all pairs of the frequency bands (Fig. 2D-F). For almost all the pairs of frequency bands (*e*.*g*., delta *vs*. theta in the hippocampus), the correlation coefficients were significantly higher than the surrogates both in the light and dark periods, but for some pairs of frequency bands, the correlation was significantly high in the light period, but not in the dark period (*e*.*g*., sniffing *vs*. slow gamma in the olfactory bulb; *P* < 1.0 × 10^−3^ (dark) and *P* = 0.98 (light), surrogate data test), and vice versa (*e*.*g*., theta *vs*. fast gamma in the hippocampus; *P* = 0.18 (dark) and *P* = 1.0 × 10^−3^ (light), surrogate data test) (Fig. 2D-F).

**Figure 2.**
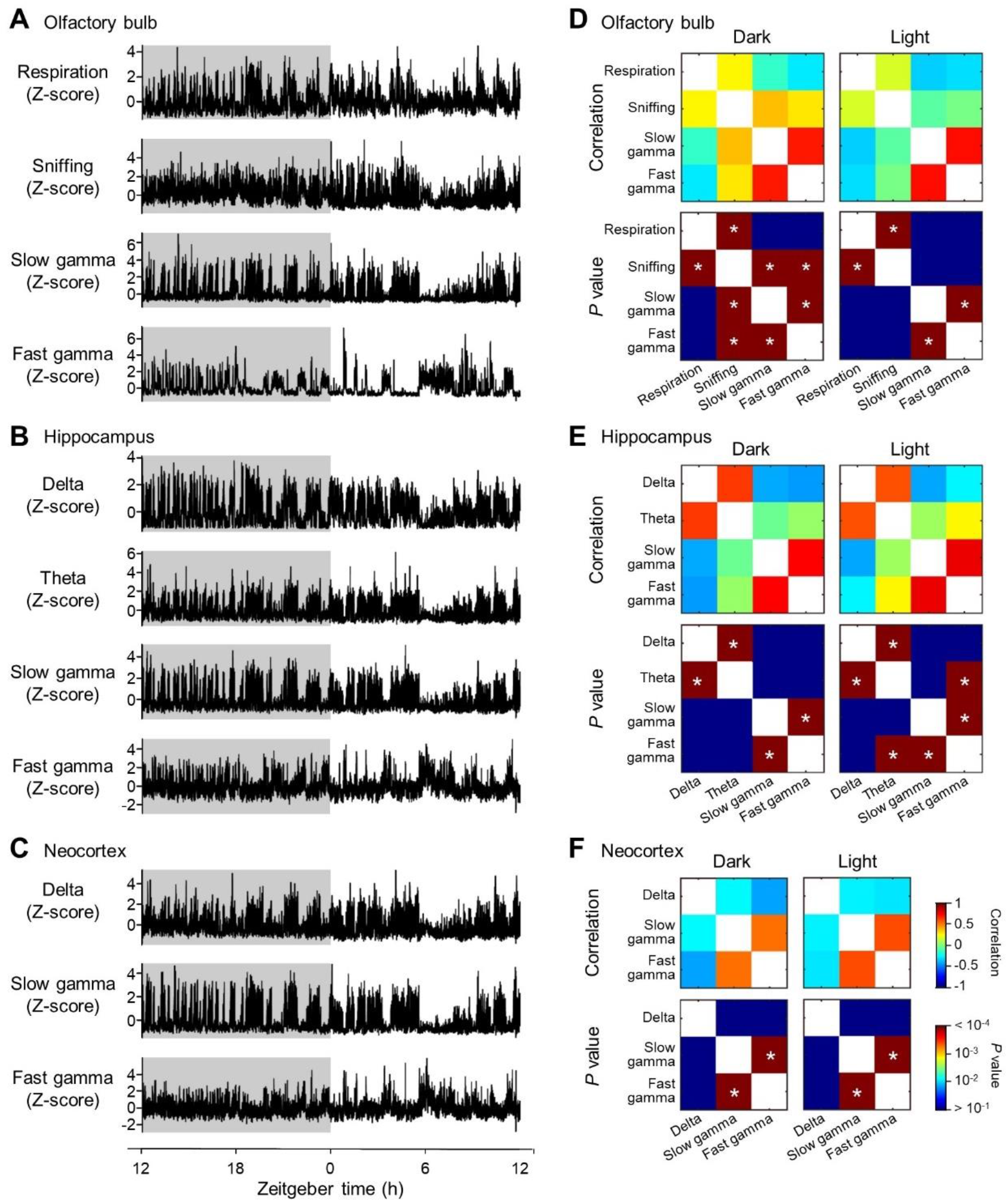
Correlation of amplitudes between pairs of bandpassed LFPs in the olfactory bulb, hippocampus, and neocortex. ***A***, Representative Z-scored amplitudes of 24 h-long olfactory bulb LFPs bandpassed at 1-4 Hz (*top* (*first*)), 6-12 Hz (*second*), 25-50 Hz (*third*), and 65-130 Hz (*fourth*), each of which represents respiration, sniffing, slow gamma, and fast gamma oscillations, respectively. ***B***, Representative Z-scored amplitudes of 24 h-long hippocampal LFPs bandpassed at 1-4 Hz (*top* (*first*)), 6-12 Hz (*second*), 25-50 Hz (*third*), and 65-130 Hz (*fourth*), each of which represents delta, theta, slow gamma, and fast gamma oscillations, respectively. ***C***, Representative Z-scored amplitudes of 24 h-long neocortical LFPs bandpassed at 1-4 Hz (*top* (*first*)), 25-50 Hz (*second*), and 65-130 Hz (*third*), each of which represents delta, slow gamma, and fast gamma oscillations, respectively. ***D***, Representative pseudocolored matrices that signify correlation coefficients of Z-scored amplitudes between pairs of bandpassed LFPs in the dark (*northwest*) and light (*northeast*) periods are shown. Hot and cold colors indicate high and low correlation coefficients, respectively (described in ***F***). Representative pseudocolored matrices that represent *P* values of corresponding correlation coefficients (mentioned above) in the dark (*southwest*) and light (*southeast*) periods are also shown. Hot and cold colors indicate high and low *P* values, respectively (described in ***F***). ***E***, Same as ***D***, but for the hippocampus. ***F***, Same as ***D***, but for the neocortex.

Since we recorded LFPs in multiple sites of each region (*i*.*e*., the olfactory bulb, hippocampus, and neocortex), we repeated these correlation analyses for all recording sites of five animals to investigate whether the number of locations where the LFP power correlations between frequency band pairs were significantly high was different between dark and light periods (Fig. 3). We found the high correlation between sniffing and slow gamma frequency powers in significantly more sites in the olfactory bulb in the dark period than the light period (*P* = 4.4 × 10^−2^, χ^2^ = 4.1, Yates’ chi-squared test), but did not for the other frequency band pairs in the olfactory bulb (Fig. 3A). We also did not any differences between the dark and light periods in the number of locations that possess high power correlations between any frequency band pairs in the hippocampus or neocortex (Fig. 3B, C).

**Figure 3.**
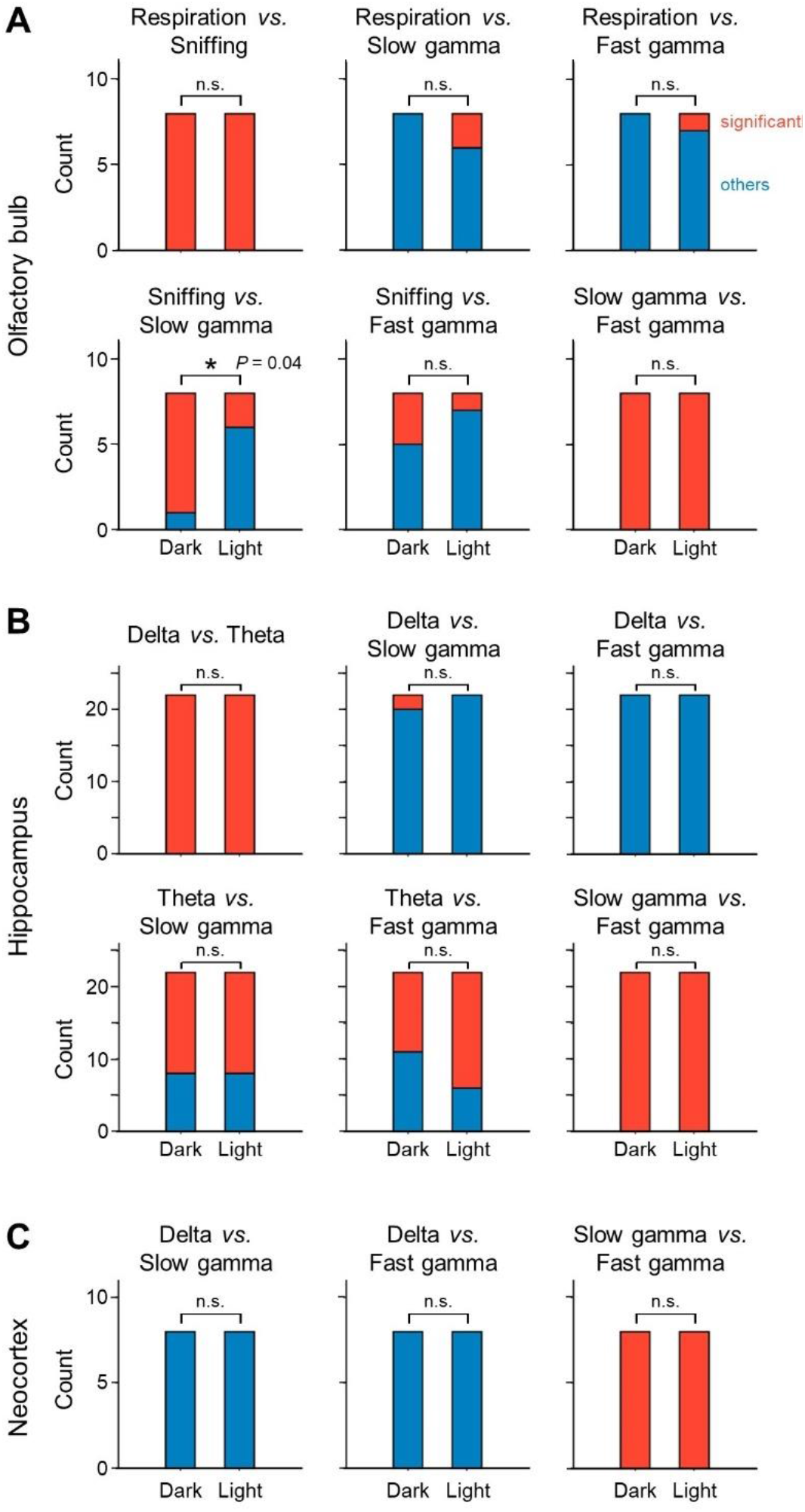
The number of recording sites where correlation between bandpassed LFPs is high in the dark and light periods. ***A***, The number of recording sites where correlation coefficients between pairs of bandpassed olfactory bulb LFPs are significantly high in the dark and light periods. ***B***, Same as ***A***, but for the hippocampus. ***C***, Same as ***A***, but for the neocortex.

## 4. Discussion

In this study, we monitored LFPs in the olfactory bulb, hippocampus, and neocortex of mice for over 24 h, enabling us to observe fluctuations in the power of each frequency band during a natural sleep-wake cycle. We found that the correlation of oscillatory powers between specific pairs of frequency bands was significantly large in the three brain regions in the dark period. In particular, the high correlation of powers between fast sniffing and slow gamma in the olfactory bulb was observed in more sites in the dark period than in the light period.

Consistent with the previous demonstration that slow gamma oscillations in the olfactory bulb is contributed by the activity of mitral cells [18], we found the high correlation between LFPs bandpassed at the sniffing and slow gamma frequency ranges. In particular, this high correlation is observed in more sites of the olfactory bulb in dark period than light period. The mechanism underlying this dark period-specific high correlation has yet to be empirically investigated, but we assume that neuromodulatory system would be responsible. Gamma oscillations in the olfactory bulb are generated by the interaction between mitral and granule cells [15]. The granule cell activity could be directly modulated by centrifugal inputs from neuromodulatory fibers that release noradrenaline, serotonin, and acetylcholine [28,29]. A previous study demonstrated that the amount of acetylcholine released in the somatosensory cortex differs between the light and dark periods [30]; thus, if the amount of acetylcholine release was fluctuated in the olfactory bulb in response to the light/dark cycle, such state-dependent fluctuation of neuromodulator levels might affect the olfactory bulb gamma oscillations, leading to fewer sites that possess high correlations between the sniffing and slow gamma oscillations in the light period.

High correlations between sniffing and gamma oscillations may indicate functional significance in that both rhythms contribute to neural coding. The olfactory system and hippocampus share electrophysiological features regarding theta rhythm (*i*.*e*., 4-12 Hz; theta oscillations (hippocampus) *vs*. sniffing (olfactory system)) and gamma rhythm (30-100 Hz); gamma oscillations are often nested in a theta-oscillatory cycle [31,32]. This phenomenon of nested theta-gamma oscillations is suggested to ‘pack’ the sequential firing of neuronal ensembles to represent a sequence of discrete items in memory [33]. In this regard, the theta cycle (*i*.*e*., sniffing for the olfactory system) includes the whole memory storage, whereas each gamma rhythm conveys an individual item held in the memory. The high correlations between sniffing and slow gamma oscillations could be attributed to the nested oscillation code. In particular, the current finding that the correlations between sniffing and gamma are high in the dark period suggests that the nested oscillation code is used more preferentially in mice deprived of light. Long-term large-scale extracellular single-unit and field recordings from the olfactory bulb of unrestrained animals will enable us to precisely unveil the coding scheme of olfactory information processing.

## Acknowledgments

The authors thank the laboratory members for helpful discussions, valuable comments, and technical support.

## Author contribution

Ryosuke Mochizuki-Koike:

- Data analysis; Writing - original draft Mami Okada:
- Data curation; Data analysis Yuji Ikegaya:
- Funding acquisition; Conceptualization; Supervision Nobuyoshi Matsumoto:
- Data analysis; Funding acquisition; Conceptualization; Supervision

